# Cytoplasmic fluidization triggers breaking spore dormancy in fission yeast

**DOI:** 10.1101/2023.09.27.559686

**Authors:** Keiichiro Sakai, Yohei Kondo, Yuhei Goto, Kazuhiro Aoki

**Author notes:** These authors contributed equally to this work (Keiichiro Sakai, Yohei Kondo).

## Abstract

The cytoplasm is a complex, crowded environment that influences myriad cellular processes including protein folding and metabolic reactions. Recent studies have suggested that changes in the biophysical properties of the cytoplasm play a key role in cellular homeostasis and adaptation. However, it still remains unclear how cells control their cytoplasmic properties in response to environmental cues. Here, we used fission yeast spores as a model system of dormant cells to elucidate the mechanisms underlying regulation of the cytoplasmic properties. By tracking fluorescent tracer particles, we found that particle mobility decreased in spores compared to vegetative cells, and rapidly increased at the onset of dormancy breaking upon glucose addition. This cytoplasmic fluidization depended on glucose sensing via the cAMP-PKA pathway. PKA activation led to trehalose degradation through trehalase Ntp1, thereby increasing particle mobility as the amount of trehalose decreased. In contrast, the rapid cytoplasmic fluidization did not require *de novo* protein synthesis, cytoskeletal dynamics, or cell volume increase. Furthermore, the measurement of diffusion coefficients with tracer particles of different sizes suggests that the spore cytoplasm impedes the movement of larger protein complexes (40–150 nm) such as ribosomes, while allowing free diffusion of smaller molecules (∼3 nm) such as second messengers and signaling proteins. Our experiments have thus uncovered a series of signaling events that enable cells to quickly fluidize the cytoplasm at the onset of dormancy breaking.

**Significance statement:** Cellular processes are influenced by the biophysical properties of the cytoplasm such as crowding and viscoelasticity. Although it has been suggested that cells tune the cytoplasmic properties in response to environmental changes, the molecular mechanisms remain unclear. Here, we used the dormant fission yeast spores and uncovered signaling pathways that facilitate cytoplasmic fluidization during dormancy breaking. Furthermore, we tracked the mobility of intracellular tracer particles, and found that the spore cytoplasm impedes the mobility of larger protein complexes, while allowing free diffusion of smaller molecules. These results suggest that small signaling proteins can diffuse relatively freely in the spore cytoplasm and have the ability to transmit dormancy breaking signals, while the motion of large complexes, such as ribosomes, is restricted.

## Introduction

The cytoplasm is a crowded environment that is densely packed with macromolecules (e.g., proteins, nucleic acids, lipids) and organelles. The crowded environments decrease the mobility of molecules, and inhibit diffusion-limited reactions by reducing the encounter rates of molecules; on the other hand, the crowded conditions can also promote intermolecular assembly through entropic effects, and facilitate reaction-limited reactions. Furthermore, accumulating evidence has shown that the cytoplasm is not a simple crowded milieu, but rather displays a complex porous structure (1–3), which is not reproduced by *in vitro* assays such as BSA solution (4, 5). In addition, there is rapidly-growing interest in the physiological significance of cytoplasmic viscoelasticity and pH (6, 7). The cytoplasmic structures and physico-chemistry influence many different biochemical reactions and cellular organization (8, 9), including microtubule dynamics (10), protein production (11), phase separation (12), and kinase reactions (13–15).

The biophysical properties, such as the structures and physico-chemistry, of the cytoplasm are disturbed by environmental changes. For example, hyper-osmotic stress enhances molecular crowding through dehydration, which inhibits macromolecular movement (10, 12, 16, 17). Energy depletion and nutrient starvation also cause reduced motion of macromolecules in the cytoplasm (7, 18–21). However, to adapt to environmental changes, cells maintain homeostasis by autonomously regulating the biophysical properties of the cytoplasm. A recent study has shown that mammalian cells possess a molecular crowding sensor, with-no-lysine kinase 1 (WNK1), which leads to cell volume recovery and reduced crowding in response to hyperosmotic stress (22). As another example, upon temperature increase, budding yeast cells synthesize trehalose and glycogen to increase cytoplasmic viscosity, which counteracts the increase in protein diffusivity (23).

Dormancy is a cellular state, in which the metabolic activity is decreased and the cell cycle is reversibly arrested under unfavorable conditions (7, 24). Those characteristics link to the cytoplasmic properties of dormant cells with reduced fluidity and molecular mobility. For example, it is known that the cytoplasms of dormant cells show a solid or glass-like properties due to their reduced water content; such characteristic cytoplasms are observed in diverse species such as bacterial spores (25, 26), fungal spores (27), plant seeds (28), tardigrades (29), and sleeping chironomids (30). Recently, micro-rheological measurements revealed that macromolecular mobility is restricted inside the cytoplasm of dormant cells (7, 19, 24). However, it remains unclear what molecular mechanisms are involved in the reversible change in cytoplasmic fluidity, and which and how intracellular signaling pathways govern these processes. In particular, since the cytoplasmic properties in a dormant state suppress intracellular diffusion and reaction, it is puzzling how dormant cells can still achieve a rapid resumption of cell growth when environmental conditions improve.

Here, we demonstrate that the glucose-sensing pathway triggers cytoplasmic fluidization during dormancy breaking of spores in the fission yeast *Schizosaccharomyces pombe*. The fission yeast enters a dormant state by forming spores under nutrient starvation, and glucose refeeding breaks the dormancy, which is called germination. By using micro-rheological tracer particles, we show that the cytoplasmic fluidity of spores is limited compared to that of vegetative cells. The solidified cytoplasm in spores rapidly fluidizes upon glucose stimulation. This fluidization process is dependent on the degradation of trehalose, which is regulated by the cAMP-PKA pathway. Based on our findings using tracer particles of different sizes, we propose the existence of a dormancy-specific intracellular milieu, which impedes diffusion of large protein complexes such as ribosomes, while allowing relatively free diffusion of small signaling proteins.

## Results

### Fission yeast spores restrict the movement of particles with a diameter of 40 nm

In this study, we adopted the fission yeast *S. pombe* spore as a model system for studying the molecular mechanisms underlying the change in cytoplasmic fluidity during dormancy breaking. To evaluate the biophysical properties in the cytoplasm of the fission yeast spore, we first compared the motion of foreign tracer particles between vegetative cells and spores. Due to the rigid cell wall, it is technically challenging to inject tracer particles into fission yeast cells (31). To overcome this limitation, we used genetically encoded multimeric nanoparticles (GEMs), which self-assemble into a spherical particle with a diameter of 40 nm (hereafter referred to as 40-nm GEMs) (Fig. 1A) (12). The gene encoding 40-nm GEMs fused to a fluorescent protein, T-Sapphire, was introduced into the fission yeast, resulting in the formation of bright tracer particles in both vegetative cells and spores (Fig. 1B, Movie S1). The GEM particles were tracked to obtain trajectories and estimate the effective diffusion coefficient (*D*_eff_) from the mean-squared displacement (MSD) curves (see Materials and Methods). The median effective diffusion coefficient was 0.54 μm^2^/s in vegetative cells (Fig. 1C and D), which was comparable to the diffusion coefficient values in the previous studies using 40-nm GEMs in fission yeast vegetative cells (10, 32–34). As we expected, the median effective diffusion coefficient of 0.025 μm^2^/s in spores indicates that the GEMs in spores were approximately 20 times less mobile compared with those in vegetative cells (Fig. 1C and D). In addition, using the ensemble-averaged MSD curves of vegetative cells and spores (inset, Fig. 1C), we computed the anomalous exponent to be ∼0.9 in vegetative cells and ∼0.75 in spores. This means that the spore cytoplasm made the GEM motion strongly subdiffusive; in other words, the GEM motion in spores was hindered by, for example, the enhanced macromolecular crowding and/or higher cytoplasmic viscoelasticity. After the spores were purified and stored at 4°C, GEM mobility was measured over time. GEM mobility decreased slightly as the spores were stored at 4°C (Fig. S1A and B), indicating that the GEM mobility does not substantially change during the storage of spores at 4°C. These results show, for the first time, that the cytoplasmic fluidity in fission yeast spores is strongly restricted compared to that in vegetative cells.

**Fig 1.**
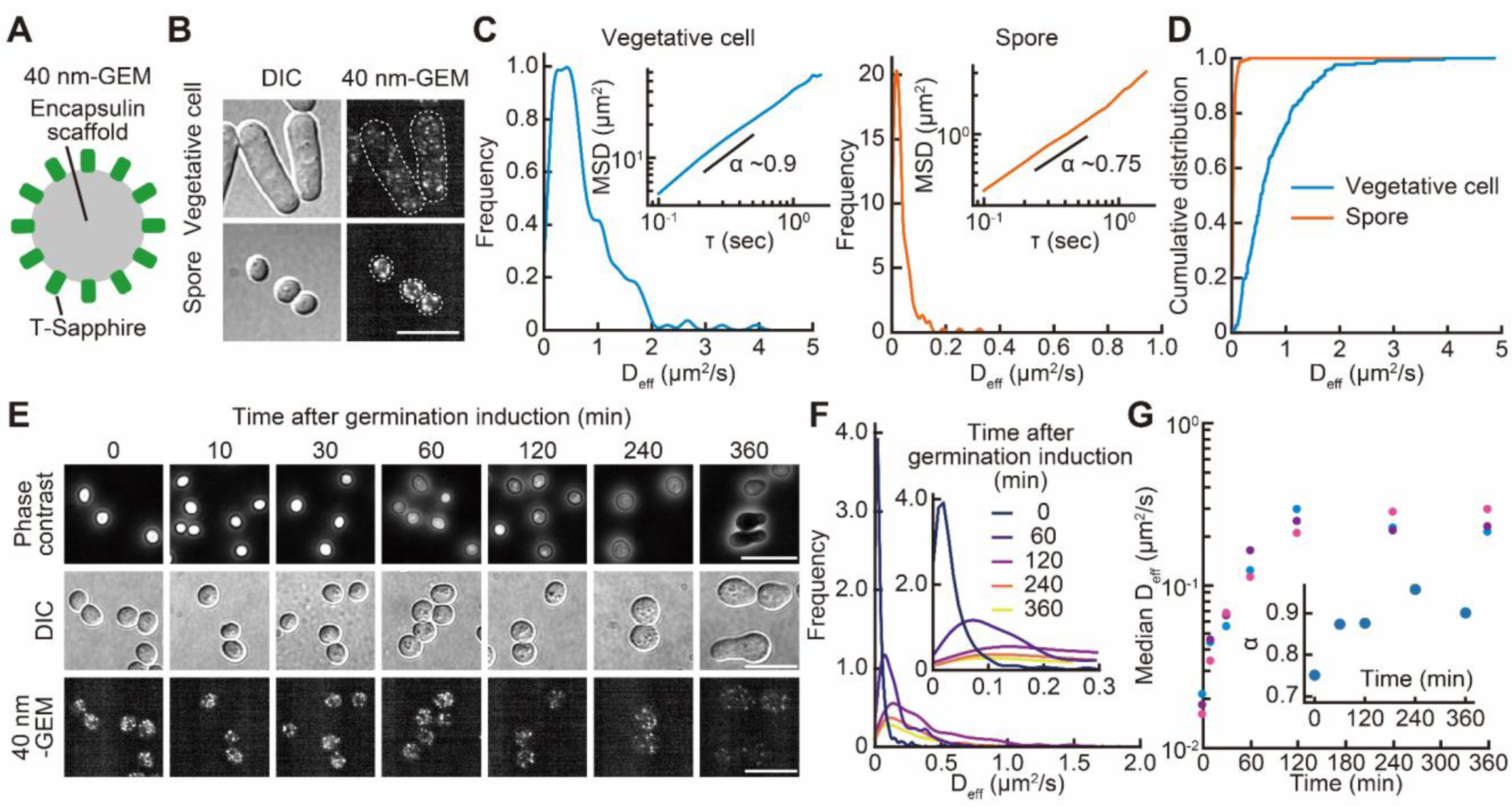
The mobility of 40 nm particles is limited in fission yeast spores but increases rapidly during germination. (A) A schematic illustration of genetically encoded nanoparticles (GEMs) with a diameter of 40 nm (40-nm GEMs). (B) Representative differential interference contrast (DIC) (left) and spinning disk confocal fluorescence images (right) of vegetative cells (upper) and spores (lower) of fission yeast expressing 40-nm GEMs. Scale bar, 10 μm. (C) Distribution of effective diffusion coefficients (*D*_eff_) for 40-nm GEMs in vegetative cells (left) (n = 218) and spores (right) (n = 221). The inset indicates the ensemble-averaged MSD curves of 40-nm GEMs for vegetative cells (left) and spores (right). (D) Cumulative distribution function of *D*_eff_ for 40-nm GEMs in vegetative cells (blue) and spores (orange). (E) Representative phase contrast images (upper) of cells expressing 40-nm GEMs at the indicated time after germination initiation upon glucose refeeding. Also shown are DIC (middle) and confocal fluorescence images (lower) under the same condition. Scale bar, 10 μm. (F) Distribution of *D*_eff_ for 40-nm GEMs at the indicated time after germination initiation (0 min, n = 801; 60 min, n = 925; 120 min, n = 1023; 240 min, n = 533; 360 min, n = 416). The inset shows the distribution of *D*_eff_ for 40-nm GEMs in the range of low values. (G) Median *D*_eff_ for 40-nm GEMs during germination. The results of three independent experiments are plotted with different colored dots (n > 300 for each condition). The inset indicates the representative anomalous exponent α during germination.

### The mobility of 40 nm particles increases during the initial stage of spore germination

Next, we examined when the reduced 40-nm GEMs mobility in spores recovered during dormancy breaking. For induction of germination, spores were transferred to glucose-rich medium. Consistent with previous reports (35–39), phase contrast microscopy indicated that the spore cytoplasm underwent a bright-to-dark transition approximately 1 hr after induction, followed by spore swelling and germ tube elongation after 4–6 hr (Fig. 1E). We tracked the GEM particles and found that the effective diffusion coefficient rapidly increased by 7-fold within the first hour after induction and subsequently reached the same level as that in vegetative cells (Fig. 1F and G, Movie S2). The subdiffusive anomalous exponent also showed the rapid recovery (inset, Fig. 1G and Fig. S2). Interestingly, the GEM mobility began to increase before the bright-to-dark transition under phase contrast microscopy, which is traditionally the earliest hallmark of germination onset (35) and signaling dynamics (37, 40).

### Glucose-sensing via the cAMP-PKA pathway is essential for the increase in the mobility of 40 nm particles during germination

To investigate the molecular mechanism underlying the observed increase in 40-nm GEMs mobility in the early phase of germination (Fig. 1F and G), we first focused on glucose-sensing via the cyclic adenosine monophosphate-protein kinase A (cAMP-PKA) pathway (Fig. 2A) (41–43). Glucose is recognized by a G-protein coupled receptor, Git3, at the plasma membrane, thereby activating the adenylate cyclase, Cyr1 (Fig. 2A). The activated Cyr1 produces cAMP, which binds to the PKA regulatory subunit, Cgs1, and finally activates the PKA catalytic subunit, Pka1(Fig. 2A). This pathway is known to play a critical role in the initiation of germination in fission yeast spores (35). Indeed, in the strains *pka1*Δ and *cyr1*Δ, these spores did not exhibit any bright-to-dark transition (Fig. 2B) or germ tube elongation (Fig. 2C) even 15 hr after the addition of glucose. We therefore examined whether the cAMP-PKA pathway is required for the increase in the GEM mobility, which precedes the canonical early event of germination such as bright-to-dark transition and germ tube elongation. We found that the GEM mobility remained unchanged in mutant strains lacking cAMP-PKA pathway components (*pka1*, *cyr1*, or *git3*) up to 6 hr after germination induction (Fig. 2D). This result indicates that the activation of the cAMP-PKA pathway in response to glucose is necessary for the increase in the GEM mobility during germination.

**Fig 2.**
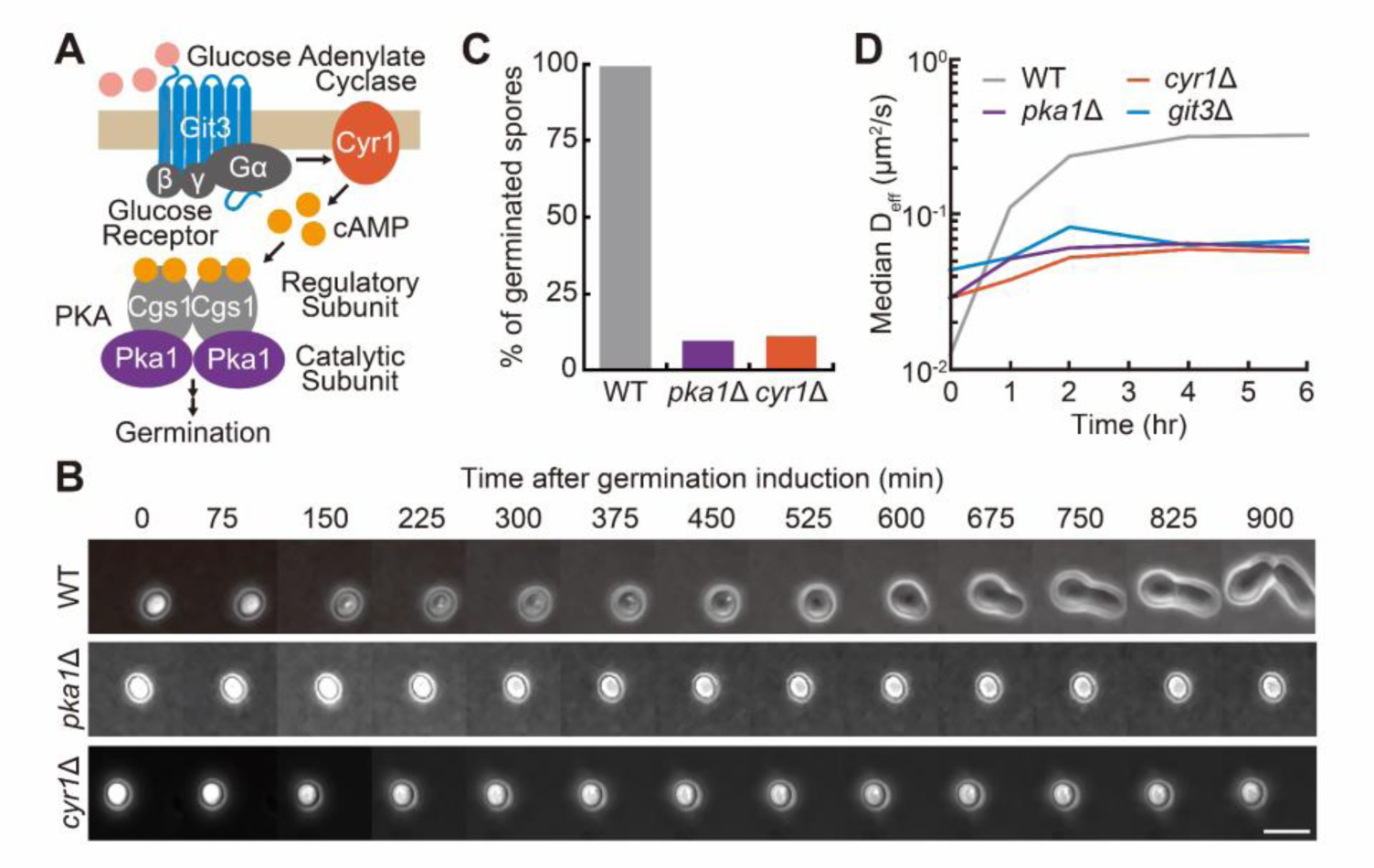
The cAMP-PKA pathway is required for the increase in the mobility of 40 nm particles during spore germination. (A) A schematic illustration of the cAMP-PKA pathway upon glucose stimulation in fission yeast. (B) Representative phase contrast images of wild-type (WT, upper), *pka1*Δ (middle), and *cyr1*Δ (lower) spores at the indicated time after germination induction. Scale bar, 5 μm. (C) Percentage of germinated spores showing an elongated germ tube at 15 hr (900 min) after the glucose addition (WT, n = 147 cells; *pka1*Δ, n = 356 cells; *cyr1*Δ, n = 163 cells). (D) Median *D*_eff_ of 40-nm GEMs during germination in the wild-type (WT), *pka1*Δ, *cyr1*Δ, and *git3*Δ strains (n > 250 for each condition).

### Germination-induced changes in particle assembly, cellular volume, protein synthesis, and cytoskeletal dynamics have no impact on the increase in mobility of 40 nm particles

In addition to the cAMP-PKA pathway, we explored other possible mechanisms affecting the mobility of 40-nm GEMs during germination. We first examined the change in particle assembly during the germination. The fluorescence intensity of each single particle appeared to decrease in spores as the germination proceeded (Fig. 1E), which might reflect the change in multimerization and/or size of the GEM particles. To examine the effects on the increase in the 40-nm GEM mobility, we quantified the fluorescence intensity of a single GEM particle at different time points (0, 1, 2, 4, 6 hr) from germination in spores and in vegetative cells (Fig. 3A). The distributions of fluorescence intensity of 40-nm GEMs in germinating spores and vegetative cells showed unimodal distribution, and their peaks shifted leftward as the germination proceeded (Fig. 3A). Although 6 hr after germination induction the average fluorescence intensity of particles was 1.5-fold less than before induction, the fluorescence intensity did not change within the first 2 hr of germination induction (Fig. 3A and B). Importantly, the 40-nm GEM mobility increased significantly within an hour of germination induction (Fig. 1F and G). From these results, we concluded that the rapid increase in the 40-nm GEM mobility was not attributable to the change in particle assembly and/or size during germination.

**Fig 3.**
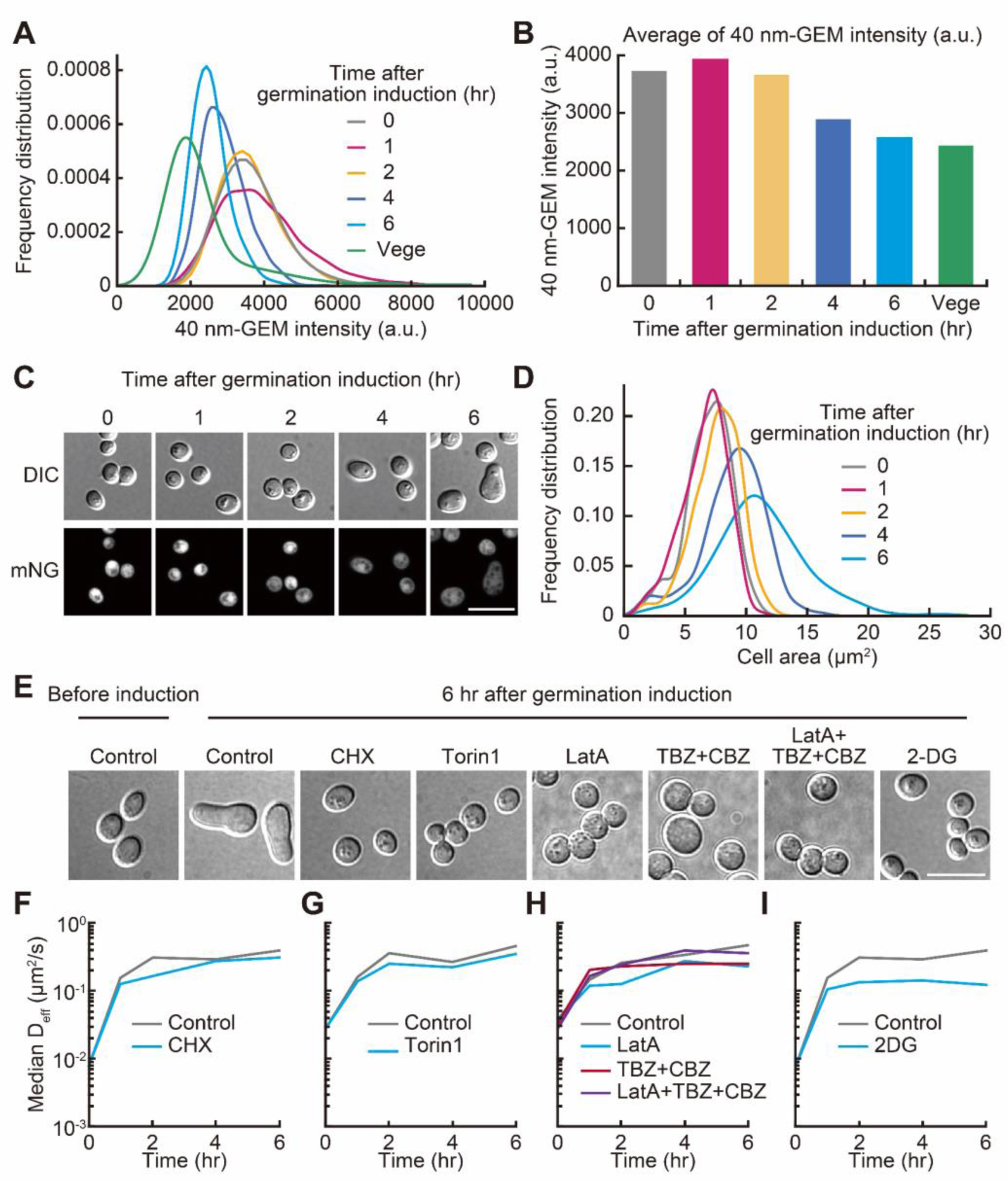
Effect of changes in particle assembly, cellular volume, protein synthesis, cytoskeletal dynamics and glycolysis on 40 nm particle mobility during germination. (A) Distribution of fluorescence intensity for 40-nm GEMs of spores at the indicated time after germination induction and vegetative cells (Vege) (n > 190 trajectories for all conditions). (B) Average values of the GEM fluorescence intensity of spores at the indicated time after germination induction and vegetative cells. The values were calculated using the data in panel A. (C) Representative DIC (upper) and confocal fluorescence images (lower) of cells expressing mNeonGreen (mNG) under the constitutive promoter *Padh1* during germination. Scale bar, 10 μm. (D) Distribution of cytoplasmic areas of spores at the indicated time after germination induction. Cytoplasmic areas were measured using images of cells expressing mNG as shown in panel C (n > 500 cells at each time point). (E–I) Effect of various types of inhibitors on germination progression and 40-nm GEMs mobility. (E) Representative DIC images of germinating spores at 0 hr and 6 hr after glucose addition in the absence or presence of cycloheximide (CHX), Torin1, latrunculin A (LatA), thiabendazole (TBZ), carbendazim (CBZ), and 2-deoxyglucose (2-DG). Scale bar, 10 μm. (F–I) Median *D*_eff_ of 40-nm GEMs during germination in the presence of CHX (F), Torin1 (G), LatA, TBZ, CBZ (H), and 2-DG (I) (n > 170 for each condition).

Prior works have shown that a reduction in cellular volume decreases macromolecular mobility under hyper-osmotic stress and nutrient starvation (10, 16, 18). Indeed, the cellular volume increases as the germination proceeds (Fig. 1E) (35, 37). We therefore asked whether the increase in cellular volume contributed to the increase in the 40-nm GEM mobility after germination induction. For simplicity, we quantified cytoplasmic areas by using spores expressing mNeonGreen (Fig. 3C). The cytoplasmic area of spores did not change within 2 hr after germination induction, but increased thereafter (Fig. 3D). Thus, these results strongly suggest that the increase in cytoplasmic volume during germination does not cause the increase in the 40-nm GEM mobility in the early stage of germination.

Finally, we explored the effects of protein synthesis or cytoskeleton on the 40-nm GEM mobility by using small chemical inhibitors. Treatment with a translation inhibitor, cycloheximide (CHX), or a TOR inhibitor, Torin1, did not change the effective diffusion coefficients of 40-nm GEMs during germination (Fig. 3E–G). Meanwhile, both CHX and Torin1 blocked the swelling and germ tube formation caused by germination induction (Fig. 3E), indicating that protein synthesis is necessary for the processes of swelling and germ tube formation, but not for the increase in the 40-nm GEM mobility in the earlier stage of germination. We next asked whether the cytoskeletal structure and/or dynamics influence the 40-nm GEM mobility. Intracellular particle diffusion is known to be affected by cytoskeletons and molecular motors (12, 44). To examine these effects, we treated cells with an actin depolymerizer, latrunculin A (LatA), and/or microtubule-destabilizing agents, thiabendazole (TBZ) and carbendazim (CBZ), but none of these compounds had any impact on the 40-nm GEM mobility (Fig. 3H). The germ tube formation was inhibited by treatment with these drugs (Fig. 3E). These results demonstrate that the cytoskeletal changes make a negligible contribution to the increase in the 40-nm GEM mobility during germination.

### ATP production via glycolysis contributes to an increase in 40 nm particle mobility after 1 hr of germination

Previous studies have shown that ATP depletion drastically reduces tracer particle diffusion in bacterial cells (7, 19, 44), yeast cells (7, 45), and mammalian cells (44). We therefore investigated whether *de novo* ATP synthesis affords the 40-nm GEM mobility in the germination process. Previous studies reported that a glycolysis inhibitor decreased the ATP level in vegetative cells of *S. pombe* (46, 47). In light of these reports, we treated spores with an inhibitor of glycolysis, 2-deoxyglucose (2-DG), which resulted in the inhibition of swelling and germ tube formation at 6 hr after germination induction (Fig. 3E). This result indicates that ATP synthesis via glycolysis is required for the germination progression. Interestingly, the 40-nm GEM mobility increased within an hour after germination induction even in the presence of 2-DG, but remained at a constant level thereafter (Fig. 3I). From these results, we found that the 40-nm GEM mobility was independent of ATP synthesis via the glycolysis immediately after germination initiation but was dependent on it from 1 hr after germination initiation. However, the effect of glycolysis on 40-nm GEM mobility after 1 hr of germination initiation was relatively small, increasing the effective diffusion coefficient by only 2-to 3-fold.

### Trehalose degradation downstream of the cAMP-PKA pathway increases 40 nm particle mobility during germination

Among the possible mechanisms for increased 40 nm GEM mobility during germination, we focused on intracellular trehalose, a disaccharide formed by a 1,1-glycosidic bond between two glucose molecules. As previously demonstrated (48, 49), trehalose in spores accumulated nearly 1,000-fold compared to the level in vegetative cells (Fig. 4A). We roughly estimated that the trehalose concentration was approximately 5% w/v in a fission yeast spore (see Materials and Methods). Furthermore, recent studies have shown that vegetative cells of budding yeast respond to heat shock by accumulating trehalose, which increases cytoplasmic viscosity and thus maintains a constant protein diffusion rate (23). Therefore, we hypothesized that trehalose accumulation causes a decrease in cytoplasmic viscosity in spores and that trehalose degradation triggers the cytoplasmic fluidization upon germination.

**Fig 4.**
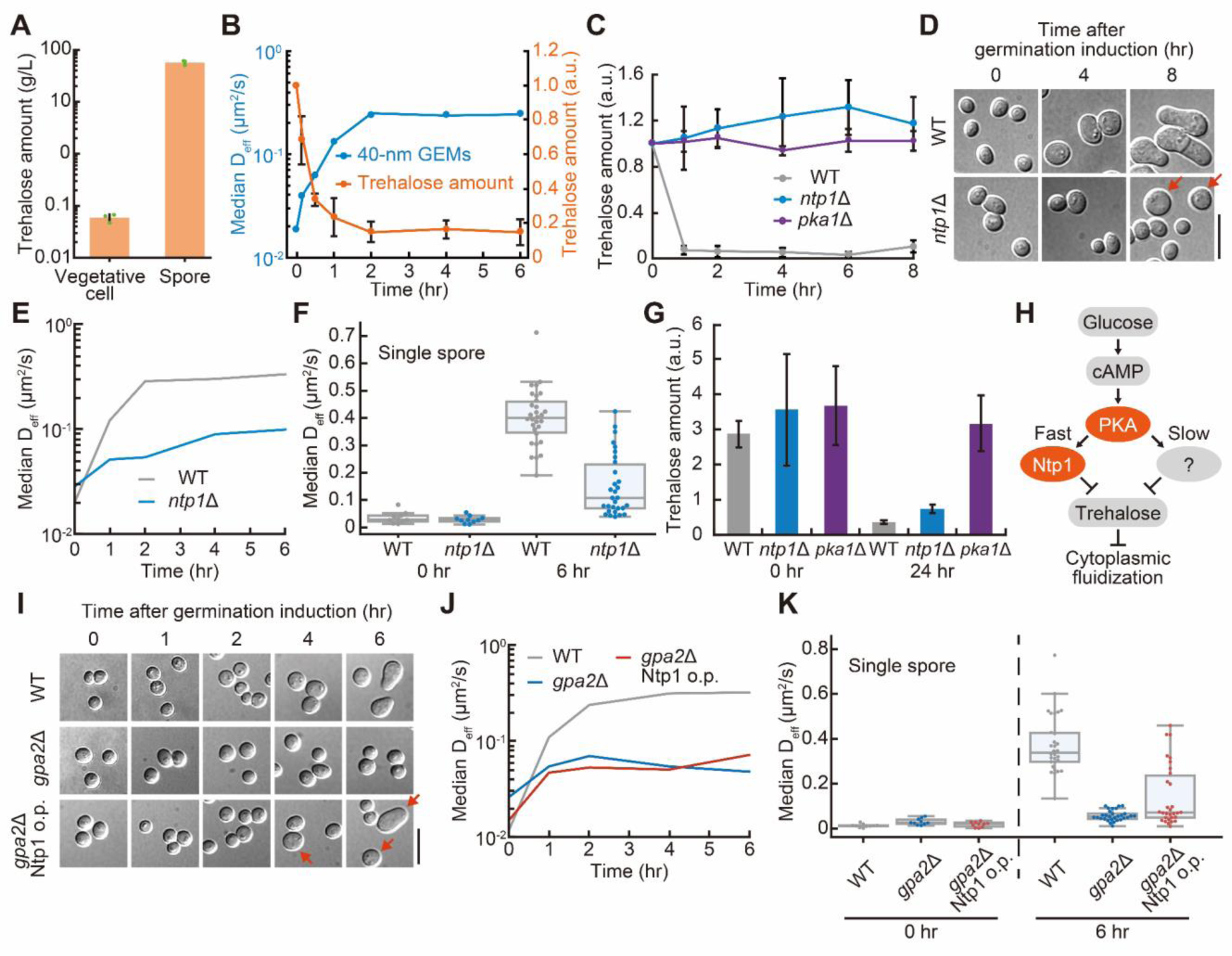
Trehalose degradation through the cAMP-PKA pathway is required for the increase in the mobility of 40 nm particles. (A) Quantification of the trehalose amount in vegetative cells and spores. Mean values were calculated from three independent experiments, each indicated by a green dot. Error bars represent SD. (B) Time-courses of 40-nm GEM mobility and trehalose amount at the indicated time after germination induction. A blue line indicates the 40-nm GEM mobility, which was calculated as the mean of the three median *D*_eff_ values in Fig. 1G. An orange line indicates the trehalose amount, which was the mean from three independent experiments. Error bars represent SD. (C) Trehalose amount in wild-type (WT), *ntp1*Δ, and *pka1*Δ spores at the indicated time after germination induction. Each line indicates the mean value from three independent experiments. Error bars represent SD. (D) Representative DIC images of WT (upper) and *ntp1*Δ (lower) at the indicated time after germination induction. Red arrows indicate swollen spores. Scale bar, 10 μm. (E) Median *D*_eff_ of 40-nm GEMs during germination in WT and *ntp1*Δ spores (n > 250 for each condition). (F) Median *D*_eff_ of 40-nm GEMs at the single spore level for WT and *ntp1*Δ strains. Each dot represents the median *D*_eff_ of 40-nm GEMs of a single cell with a boxplot, in which the box shows the quartiles of data with the whiskers denoting the minimum and maximum except for the outliers detected by 1.5 times the interquartile range (n = 10 cells at 0 hr and n = 30 cells at 6 hr). (G) Trehalose amounts in the WT, *ntp1*Δ, and *pka1*Δ strains at 0 hr and 24 hr after germination induction. Mean values from three independent experiments are shown with error bars (SD). (H) A schematic model of the glucose sensing and trehalose degradation pathways after germination initiation upon glucose refeeding. PKA activation leads to trehalose degradation via Ntp1-dependent and -independent pathways, resulting in cytoplasmic fluidization and germination. (I) Representative DIC images of spores of the WT (upper), *gpa2*-deficient mutant (*gpa2*Δ; middle), and *gpa2*-deficient mutant expressing Ntp1 under the spore-inducible promoter *Posi2* (*gpa2*Δ Ntp1 o.p.; lower) upon germination induction. Red arrows indicate swollen spores starting to germinate. Scale bar, 10 μm. (J) Median *D*_eff_ of 40-nm GEMs during germination in the WT, *gpa2*Δ, and *gpa2*Δ Ntp1 o.p. spores (n > 250 for each condition). (K) Median *D*_eff_ of 40-nm GEMs at the single spore level for the WT, *gpa2*Δ, and *gpa2*Δ Ntp1 o.p. strains. Each dot represents the median *D*_eff_ of 40-nm GEMs of a single cell with a boxplot as in panel F (n = 10 cells at 0 hr and n = 30 cells at 6 hr).

To investigate this hypothesis, we first quantified how the trehalose amount in fission yeast spores changed during germination, and found that trehalose was rapidly degraded on the same timescale as the increase in the 40-nm GEM mobility (Fig. 4B). We then measured the trehalose amount and the GEM mobility in a mutant strain lacking *ntp1*, a trehalase gene (50). We found that *ntp1*-deleted cells (*ntp1*Δ) did not show the rapid decrease in trehalose amount within 1 hr of germination induction observed in WT cells, and their trehalose amount remained constant up to 8 hr after germination induction (Fig. 4C). Additionally, in spores lacking *pka1*, trehalose degradation was suppressed after germination induction (Fig. 4C), implying that the cAMP-PKA pathway controls the trehalose degradation pathway. We also found that the germination process was severely delayed in the *ntp1*Δ strain, which is incapable of forming a germ tube 8 hr after germination (Fig. 4D), consistent with a previous report (49). Importantly, the increase in 40-nm GEM mobility was substantially suppressed in the *ntp1*Δ strain compared to the WT after germination induction (Fig. 4E). These results suggest that Ntp1-mediated trehalose degradation is required for the cytoplasmic fluidization and rapid germination through the cAMP-PKA pathway.

We recognized that a small fraction of spores in the *ntp1*Δ strain displayed swelling, an initial feature of germination, 8 hr after germination induction (Fig. 4D, red arrows). This result could be due to the Ntp1-independent trehalose degradation in this small population of spores, resulting in the progression of germination. Indeed, the quantification of the effective diffusion coefficients of GEMs at the single spore level revealed that approximately one-third of *ntp1*Δ spores showed an increase in the 40-nm GEM mobility comparable to wild-type spores 6 hr after germination induction (Fig. 4F). Consistent with these results, the trehalose amount decreased even in *ntp1*Δ spores 24 hr after the germination induction, while it remained unchanged in *pka1*Δ spores (Fig. 4G). Taken together, these results lead us to propose the existence of at least two trehalose degradation pathways during the spore germination of fission yeast; one is Ntp1-dependent fast trehalose degradation, whereas the other is Ntp1-independent slow trehalose degradation (Fig. 4H). Both trehalose degradation pathways are controlled by the cAMP-PKA pathway (Fig. 4H). We note that the former pathway, Ntp1-dependent fast trehalose degradation, is necessary for rapid germination in response to glucose stimulation.

### Overexpression of trehalase in spores deficient in the cAMP-PKA pathway partially increases the mobility of 40 nm particles during germination

We next examined whether Ntp1 overexpression is sufficient to increase the 40-nm GEM mobility and induce germination. First, we developed a spore-specific gene expression system in fission yeast cells by using promoters whose expression was specifically induced after the completion of the sporulation process. By reanalyzing previously reported microarray data during sporulation in fission yeast cells (51), we identified the top 4 genes (*SPAC869.09*, *hry1*, *isp3*, *pdc202*) whose expression was strongly induced in spores compared to vegetative cells. The region approximately 1,000 bp upstream of those genes was used as the promoter region to express a certain gene in spores. We named these promoters *Posi1-4* (<u>P</u>romoters for <u>o</u>nly <u>s</u>pore induction), respectively. We quantified the expression levels of mNeonGreen under *Posi1-4* (Fig. S3A–D), and found that *Posi2* showed the highest ratio of the expression of mNeonGreen in spores to that in vegetative cells (Fig. S3E). Therefore, we used *Posi2* for the following experiment.

To investigate whether Ntp1 overexpression suffices to induce germination, we established and analyzed spores overexpressing Ntp1 under *Posi2*. However, the spores overexpressing Ntp1 did not show any morphological change in the absence of glucose and exhibited an accelerated increase in the 40-nm GEM mobility upon glucose stimulation. Therefore, we next overexpressed Ntp1 in spores deficient in the cAMP-PKA pathway and treated the spores with glucose in order to examine the effects of Ntp1 overexpression on glucose-induced morphological change and the 40-nm GEM mobility. Spores lacking *gpa2*, the gene encoding heterotrimeric G protein alpha subunit (Fig. 2A), which mediates the glucose receptor and the cAMP-PKA pathway, demonstrated a significant delay in swelling upon glucose stimulation (Fig. 4I), as seen in *pka1-* or *cyr1*-deleted spores (Fig. 2B and C). Meanwhile, the overexpression of Ntp1 in spores lacking *gpa2* partially rescued the glucose-induced germination, namely swelling (Fig. 4I). The 40-nm GEM mobility was also partially increased by the overexpression of Ntp1 at the population level (Fig. 4J) and at the single cell level (Fig. 4K). These results suggest that both Ntp1 and nutrient sources such as glucose in the germination induction medium are sufficient for swelling and cytoplasmic fluidization.

### The mobility of 50–150 nm particles also decreases in spores and increases at the onset of germination

To gain further insight into the spore cytoplasm, we examined the size dependency of particle diffusion, which reflects the spatial scales of cytoplasmic structures (12, 52, 53). We first observed the mobility of particles with size larger than 40-nm GEMs by using μNS, which self-assembles into particles of different sizes (50–150 nm) (19, 54). In both vegetative cells and spores, μNS were observed as particles of varying size and brightness (Fig. 5A). We then analyzed the μNS particles with fluorescence intensities lower than a certain threshold to mitigate the effects of size variability (Fig. S4A). The effective diffusion coefficients were 0.015 μm²/s and 0.00033 μm²/s in vegetative cells and spores, respectively, indicating that the mobility of μNS, similarly to that of 40-nm GEMs, was strongly restricted in spores (Fig. 5B). During the germination process, the μNS mobility was gradually increased (Fig. 5C), which is consistent with a previous report on budding yeast spores (24). Within the first 2 hr of germination, μNS mobility increased about 6-fold, followed by a gradual increase in μNS mobility. We also quantified the anomalous exponent during germination from the ensemble-averaged MSD curves (inset, Fig. 5C and Fig. S4B). Compared with the GEMs, the μNS in spores exhibited a much smaller anomalous exponent value, α < 0.2. This result indicates that, in spores, the μNS particles were almost immobilized in the time scale of observation. To provide further support for this idea, we computed velocity autocorrelation functions (VAFs) (Fig. S4C–H), which were used as a diagnostic tool to explore the underlying mechanism of intracellular subdiffusion (7, 55). The VAFs of the spore cytoplasm showed sharper negative peaks compared to those of cells 1 hr after germination induction or vegetative cells (Fig. S4D, F, and H). These results suggest that the μNS particles are locally confined and their motion is caused by their bouncing off elastic structures as they collide with them.

**Fig. 5.**
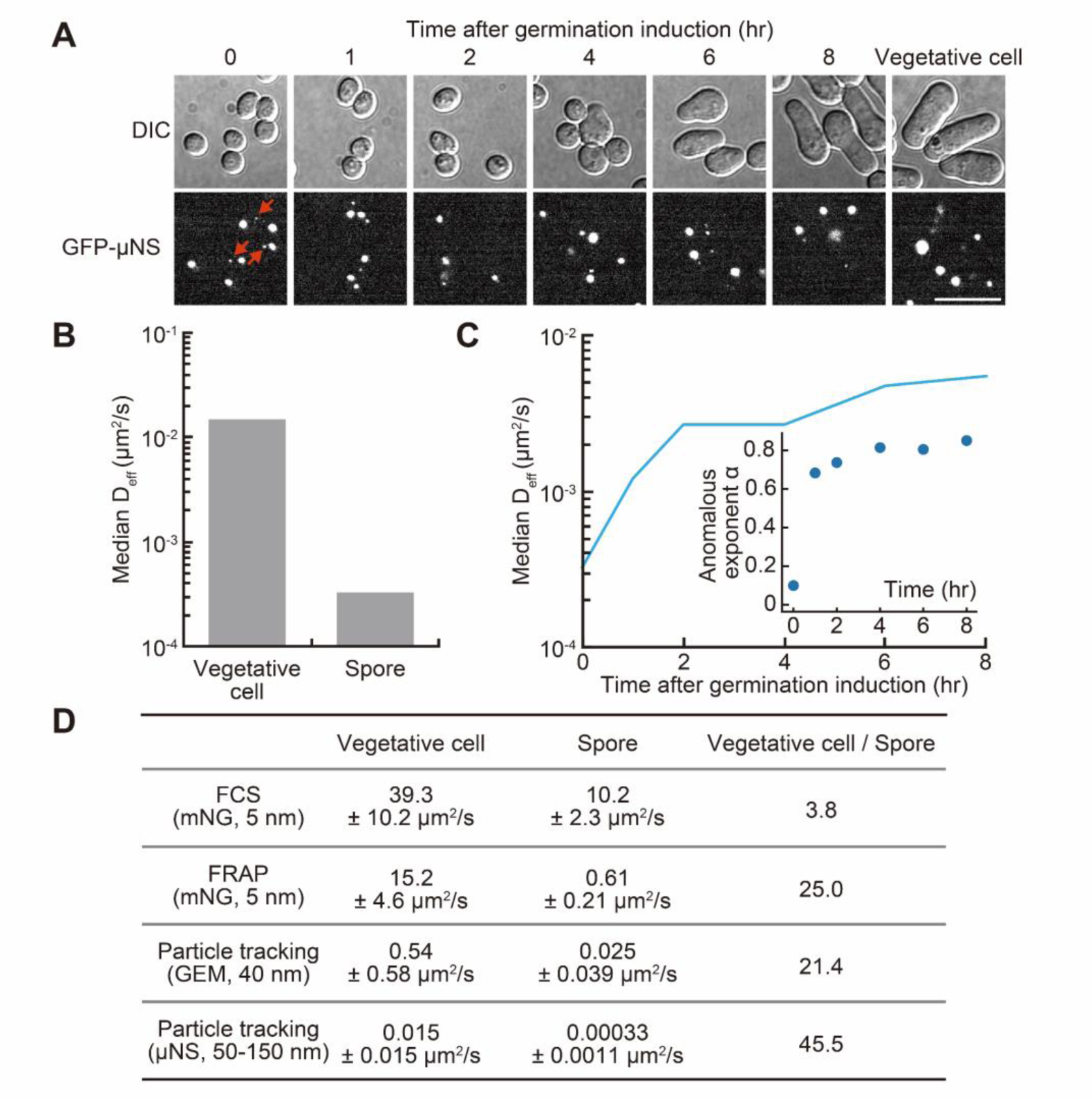
The mobility of particles of different sizes in spores and vegetative cells. (A) Representative DIC (upper) and confocal fluorescence images (lower) of fission yeast cells expressing μNS at the indicated time after germination induction. Red arrows indicate the small size particles (also see in Fig. S4A), which are used for the estimation of the effective diffusion coefficient of μNS. Scale bar, 10 μm. (B) Median effective diffusion coefficients for μNS in vegetative cells (n = 84) and spores (n = 54). (C) Median effective diffusion coefficient for μNS during germination (n > 50 for each condition). The inset indicates anomalous exponent α during germination. (D) Summary of effective diffusion coefficients for particles of various sizes in vegetative cells and spores. Vegetative cell/Spore indicates the ratio of the effective diffusion coefficients of vegetative cells to those of spores.

### Fluorescent protein-sized particles diffuse throughout the spore cytoplasm within a time frame on the order of seconds

Although the diffusion of 40-nm GEMs and μNS was drastically restricted in the spore cytoplasm (Fig. 1C and 5B), it remains unclear whether smaller proteins are also affected by the change in the biophysical properties of spores. To investigate this, we quantified the effective diffusion coefficients of the green fluorescent protein mNeonGreen (mNG, around 3 nm in size), which is smaller than 40-nm GEMs. We first observed the movement of mNG by using fluorescence correlation spectroscopy (FCS). FCS is a technique used to estimate the effective diffusion coefficient in a confocal volume (∼1 fL) by observing the fluctuation of fluorescent molecules (56–58). In the FCS analysis, we measured the time-series of mNG intensity in vegetative cells and spores of fission yeast and obtained autocorrelation functions (Fig. S5A and B). The effective diffusion coefficient of mNG in spores was about 4-fold reduced compared to that in vegetative cells (Fig. 5D and S5C). For further examination, we used fluorescence recovery after photobleaching (FRAP) to estimate the effective diffusion coefficient of mNG. Due to the tiny volume of fission yeast cells, especially the spores, a photobleached region inevitably overlaps with the cell boundary (Fig. S5D), which violates geometrical assumptions of the common fitting curve in FRAP analysis. To deal with this issue, we numerically simulated mNG diffusion by following the method of a previous FRAP study on bacterial cells (59). Then, we estimated the effective diffusion coefficient by matching the experimental and simulated FRAP time courses. In vegetative cells, the estimated effective diffusion coefficients of mNG based on FCS and FRAP were close to each other (Fig. 5D and S5E). In spores, interestingly, we found that the mNG diffusivity obtained from FRAP was much lower than that from FCS (Fig. 5D and S5E). It is not uncommon for the FRAP analysis to produce estimates of the effective diffusion coefficient of a protein that are an order of magnitude smaller than those of the FCS. Potential mechanisms underlying this discrepancy include inhomogeneous diffusivity and/or binding to immobile intracellular structures (60, 61), although determining the precise mechanism for the spore cytoplasm requires further investigation. In our present experiments, even in the FRAP analysis, mNG diffused throughout the spore cytoplasm within a time frame on the order of seconds (Fig. S5D). Based on these results, it is proposed that small signaling proteins, such as cAMP-PKA pathway and Ntp1, are relatively free to diffuse in the spore cytoplasm and have the ability to transmit dormancy breaking signals after the glucose addition, while the motion of large protein complexes is restricted.

## Discussion

Growing evidence indicates that a liquid-like cytoplasmic property turns into a solid- or glass-like state under stress conditions, and these cytoplasmic changes have been proposed to be responsible for stress resistance and dormancy breaking (7, 24). However, the molecular basis underlying such changes in the cytoplasmic properties in response to environmental stimuli is not yet fully understood. In this study, we focused on the germination of fission yeast spores as a model system of dormancy breaking. By using genetically encoded tracer particles, we found that the spore cytoplasm markedly hinders intracellular diffusion, and rapidly fluidizes upon germination induction. The cytoplasmic fluidization during germination is mediated by the degradation of intracellular trehalose. Furthermore, we revealed the signaling pathway responsible for the trehalose degradation––namely, glucose is recognized by Git3, resulting in activation of the cAMP-PKA pathway and Ntp1.

We examined the mobility of three types of particles of different sizes in the spore cytoplasm as summarized in Fig. 6. First, although the diffusivity of mNG in spores was reduced, mNG can still diffuse throughout the spore cytoplasm within a time frame on the scale of seconds (Fig. S5D). This result indicates that signaling molecules such as PKA and Ntp1 diffuse relatively freely even in the spore cytoplasm and can respond rapidly to external signals for dormancy breaking. The large discrepancy between the FRAP- and FCS-based estimates of mNG diffusivity in spores (Fig. 5D) might reflect a spore-specific cytoplasmic state, because the cytoplasm of vegetative cells did not produce such a discrepancy. An interesting possibility is the presence of immobile binding traps, which have been theoretically investigated to reconcile disparate estimates of the diffusion coefficient of Bicoid morphogen in *Drosophila* embryos (60, 62). Second, we found a marked reduction in the effective diffusion coefficients of 40-nm GEMs and strong subdiffusion of those particles in spores (Fig. 1C and G). These results mean that the mobility of large complexes such as the ribosome (∼30 nm) is severely limited in spores. Recently, *in vitro* experiments using *E. coli* and *Xenopus* cytoplasms demonstrated that an overcrowded cytoplasm can suppress the ribosome mobility and thereby the translational activity (63, 64). It is likely that the cytoplasmic biophysics of spores also suppresses the translational activity. The decrease in protein synthesis may allow spores to avoid wasting energy sources (e.g., ATP) and substrates (e.g., amino acids, nucleic acids), thus enabling their prolonged survival without nutrients. Third, the μNS particles exhibited a drastic reduction in an anomalous exponent (α ∼0.1), which means that they were almost immobilized in the spore cytoplasm (Fig. 5C). Previous studies have reported that large structures such as protein condensates (∼200–1000 nm) and organelles (∼100–1000 nm) are immobilized in energy-depleted and nutrient-starved yeast cells (7, 18–20). However, the anomalous exponent of μNS (50∼150 nm) in spores (α ∼0.1) is far smaller than that in energy-depleted yeast cells (α ∼0.6) (7). This indicates that the spore cytoplasm is an extreme environment, and probing its material properties such as viscoelasticity could pave the way for a deeper understanding of the stress resistance achieved by the spores.

**Fig 6.**
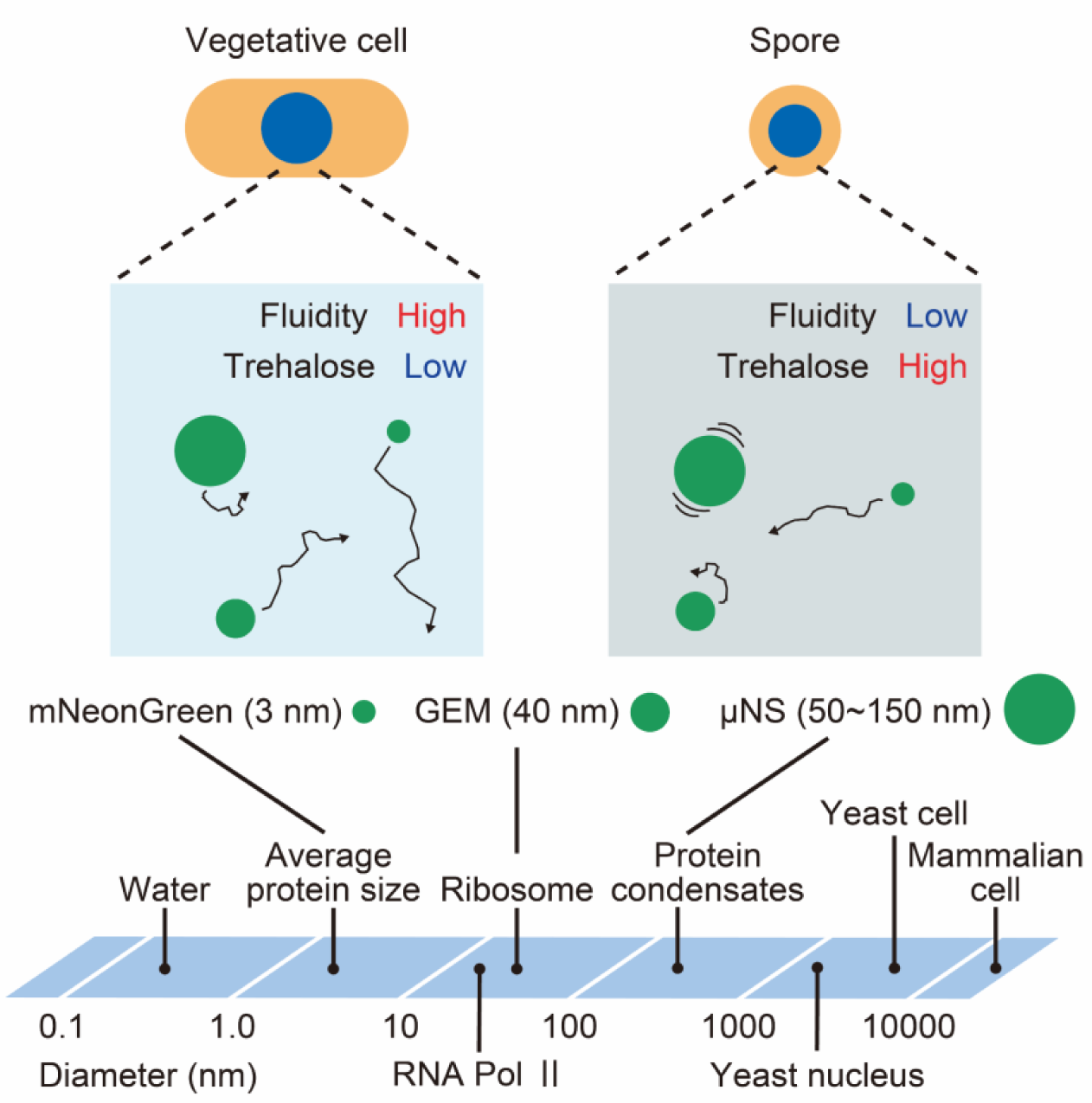
A model of cytoplasmic properties in vegetative cells and spores of fission yeast. The amount of trehalose plays a key role in regulation of the cytoplasmic properties in fission yeast cells. Vegetative cells contain only a small amount of cytoplasmic trehalose, and mNG (∼3 nm), 40-nm GEMs (∼40 nm), and μNS (∼50–150 nm) are free to diffuse in the cytoplasm. On the other hand, in spores, trehalose is highly accumulated, resulting in a decrease in the mobility of these particles. The mobility of 40-nm GEMs and μNS is strongly limited in the spore cytoplasm, while mNG are allowed to diffuse relatively freely throughout the cytoplasm.

Trehalose has been well characterized as a protein protectant and a viscosity agent *in vitro* (65). To our knowledge, however, there have been no studies on the molecular mechanism of trehalose in the regulation of cytoplasmic biophysics *in vivo* except for a few pioneering works (23, 66, 67). Our study shows that trehalose accumulates in spores at a concentration of 5% w/v (Fig. 4A), and trehalose is rapidly degraded by germination induction (Fig. 4B), resulting in a 20-fold increase in 40-nm GEM mobility. Furthermore, depletion of an enzyme that degrades trehalose, Ntp1, was shown to inhibit the increase in the 40-nm GEM mobility upon germination induction (Fig. 4E). These results imply a direct link between trehalose and cytoplasmic fluidity. However, even in a 45% trehalose solution, the diffusion rate of purified GFP was reported to be only about 2-fold lower than that in a water solution (23). Therefore, trehalose as a simple viscogen cannot explain the decrease in the 40-nm GEM mobility in fission yeast spores. We propose a model in which the trehalose forms spore-specific intracellular structures by interacting with other intracellular components. For example, it has been reported that a mixture of proteins in addition to trehalose can cooperatively increase the viscosity of a solution (68). Future studies should explore the physical chemistry of complex solutions of trehalose with proteins, lipids, and nucleic acids.

We found that glucose-triggered cAMP-PKA-Ntp1 signaling is required for cytoplasmic fluidization during germination. How is Ntp1 regulated downstream of PKA signaling in spores? There was no apparent effect of translation inhibition (CHX treatment) on the 40-nm GEM mobility during germination (Fig. 3F), suggesting that PKA and Ntp1 are already expressed in spores and their activities are regulated posttranslationally during germination. These speculations can account for the fact that spore cytoplasm could not be fully fluidized merely by overexpressing Ntp1 in the cAMP-PKA pathway-deficient spores (Fig. 4J and K). Consistent with these ideas, it has been reported that PKA-dependent phosphorylation activates trehalase Ntp1 in fission yeast and its budding yeast ortholog Nth1 (69, 70). Additionally, Ntp1 has a consensus site (S71) for PKA-dependent phosphorylation, and replacing S71 with an alanine residue results in loss of trehalase activity (71). However, it remains controversial whether PKA directly activates Ntp1 via phosphorylation (70–72). Future research should investigate the relation between PKA phosphorylation and Ntp1 trehalase activity in *in vitro* experiments.

By using foreign tracer particles, our study revealed that the biophysical properties of spores are quite different from those of vegetative cells. In addition to these exogenous particles, it would be interesting to investigate the mobility of endogenous proteins of different sizes in spores, as was recently done in a study on bacterial cells (73), because various features of proteins, such as charge and hydrophobicity, influence their mobility. In addition, the motions of intracellular particles are affected by both cytoplasmic viscoelasticity and active forces generated from molecular motors and metabolic reactions. However, in this study we relied on passive observations of the motion of probe particles, and thus we were not able to distinguish between the relative contributions of the passive and active mechanisms. A solution to this limitation would be to compare passive measurements with active microrheological measurements, in which the probe particles are controlled exogenously by optical tweezers or other means (44, 74). The prospect of applying such an approach to the spore cytoplasm is intriguing, although active microrheology experiments for yeast cells are still challenging at this time because the cell wall hinders the injection of specialized particles. Future studies will need to examine the cytoplasmic biophysics of dormant cells and how it regulates intracellular processes and functions.

## Materials and methods

### Plasmids

All plasmids used in this study are summarized in Table S1, along with Benchling links containing the plasmid sequences and maps. The nucleotide sequence of 40-nm GEM was optimized for fission yeast codon usage. To construct pMNAT2LA21-40nm-GEM_sp_opt, the DNA sequence of 40-nm GEM was subcloned into pMNAT2LA21 (75) by using conventional ligation with Ligation high Ver.2 (TOYOBO, Osaka, Japan) according to the manufacturer’s instructions. The gene was driven by a medium-strength constitutive promoter (*Padh21*) and expressed from the 2L locus (76). In a previously reported study, 40-nm GEM was developed by fusing a natural homomultimeric scaffold with the fluorescent protein T-Sapphire (12). In our present experiments, we noticed that the fluorescent protein of 40-nm GEM did not have the mutations (Q69M, C70V, V163A, S175G) that are characteristic of T-Sapphire (77), and its amino acid sequence was more similar to that of Sapphire rather than T-Sapphire. For the construction of plasmids to express mNG under the *Posi* series (pHBCO1-HA-spmNeonGreen, pHBCO2-HA-spmNeonGreen, pHBCO3-HA-spmNeonGreen, pHBCO4-HA-spmNeonGreen), a region approximately 1,000 bp upstream from the start codon in 4 genes (*SPAC869.09*, *hry1*, *isp3*, *pdc202*) was amplified by PCR and substituted for *Padh1* in pHBCA1-HA-spmNeonGreen-new by using Gibson assembly with NEBuilder HiFi DNA Assembly (New England Biolabs, Ipswich, MA). In these plasmids, mNG is expressed from the C locus (78). We used mNG optimized for fission yeast codon usage (76). To construct pHBCO2-ntp1, the DNA sequence of mNG in pHBCO2-HA-spmNeonGreen was replaced with the cDNA of *ntp1* with Gibson assembly. To construct pMNATZA1-GFP-μNS, the DNA sequence of μNS was obtained from pRS306-PHIS3-GFP-μNS (Addgene plasmid #116935; deposited by Liam Holt), and subcloned into pMNATZA1. We note that the fluorescent protein of μNS is GFPmut2, which was derived from avGFP with some mutations (S65A, V68L, S72A) (79).

### Reagents

CHX was purchased from Nacalai Tesque (#06741-91), dissolved in ethanol (100 mg/mL stock solution), and stored at −30℃. 2-DG was purchased from FUJIFILM Wako (#040-06481), dissolved in double-distilled water (DDW) (1 mM stock solution), and stored at −30℃. LatA was purchased from FUJIFILM Wako (#129-04361), dissolved in DMSO (25 mg/mL stock solution), and stored at −30℃. TBZ and CBZ were purchased from TCI (#T0830) and Sigma-Aldrich (#378674-100G), respectively, and dissolved in DMSO. The drug cocktail of TBZ and CBZ was prepared just before use. Torin1 was purchased from Selleckchem (#S2827), dissolved in DMSO (3 mM stock solution), and stored at −30℃.

### Fission yeast strains and culture conditions

All fission yeast strains used in this study are summarized in Table S2 along with their origins. Unless otherwise noted, the growth medium, sporulation medium, and other aspects of the experimental methods for fission yeast were based on those described previously (80). The transformation protocol was modified from a previously reported one (81).

### Purification of spores and induction of germination

Sporulation was induced in sporulation agar medium (SPA) at 26℃ for 3 days. SPA is composed of 1% glucose, 0.1% KH_2_PO_4_, 2% agar, and supplemental vitamins (pantothenic acid, nicotinic acid, inositol, and biotin) (82). Spores were purified by the Percoll gradient method (83) with modifications as described below. Briefly, cells on the SPA plate were suspended in 1 mL of DDW with 2% glusulase (PerkinElmer, #NEE154001EA) and digested with shaking at room temperature for 3 hr. The cells were then washed twice with 500 uL of 0.5% Triton-X and resuspended in 200 uL of 0.5% Triton-X. This suspension was layered on 1 mL of Percoll (GE Healthcare, #17089101) and centrifuged at 9,560 g for 5 min. After centrifugation, the pellet was collected and resuspended in 200 uL of 0.5% Triton-X, followed by centrifugation with Percoll using the same protocol as for the first Percoll centrifugation. The pellet was washed twice with 0.5% Triton-X, resuspended in 200 uL of Triton-X, and stored at 4℃ until used for experiments. Before germination induction, purified spores were washed twice with 1 mL of DDW and suspended in YEA medium without glucose (YEA-glucose medium) (82). Germination was induced by adding glucose to a final concentration of 2%. Spores were incubated at 32℃, and a portion of the spores was collected at each time point and used in subsequent experiments.

### Live-cell fluorescence imaging of fission yeast cells

Cells were imaged with an IX83 inverted microscope (Olympus) equipped with an sCMOS camera (ORCA-Fusion BT; Hamamatsu Photonics), an oil-immersion objective lenses (UPLXAPO 100×, NA = 1.45, WD = 0.13 mm; or UPLXAPO 60× PH, NA = 1.42, WD = 0.15 mm; Olympus), and a spinning disk confocal unit (CSU-W1; Yokogawa Electric Corporation). The excitation lasers and fluorescence filter settings were as follows: excitation laser, 488 nm for mNG, 40-nm GEM (T-Sapphire), and μNS (GFP); dichroic mirror, DM405/488/561/640 for mNG, 40-nm GEM, and μNS (Yokogawa Electric Corporation); emission filters, 525/50 for mNG, 40-nm GEM, and μNS (Yokogawa Electric Corporation). This microscope was controlled by MetaMorph software (ver. 7.10.3). For the phase-contrast imaging in Fig. 1E, cells were visualized with an BX53 upright microscope (Olympus) equipped with a digital camera (DP22; Olympus) and an air/dry objective lens (UPLFLN 40× PH, NA = 0.75, WD = 0.51 mm; Olympus).

For snapshot imaging, cells were concentrated by centrifugation at 860 g, mounted on a slide glass (thickness, 0.9–1.2 mm; Matsunami), and sealed by a glass coverslip (thickness, 0.13–0.17 mm; Matsunami). For time-lapse imaging of the germination process in Fig. 2B–C, cells were embedded in a YEA medium solidified with 2% low melting temperature agarose (Lonza, #50101) on a polylysine-coated glass bottom dish (Matsunami, #D11531H) (84).

For imaging of cells expressing 40-nm GEM, the fluorescence images were recorded every 100 milliseconds for 10 seconds in the streaming mode. The particles were tracked with the Mosaic suite, a FIJI plugin (85). In the analysis with the Mosaic suite, the following typical parameters were used: radius = 3; cutoff = 0; per/abs, variable; link range = 1; and maximum displacement = 6 pixels, assuming Brownian dynamics. Particles with tracks shorter than 10 frames were excluded from further analysis.

To quantify the intensity of particles in Fig. 3A–B, the parameter of radius in Mosaic suite was fixed at 3, and the mean intensities of each trajectory were calculated and used as a proxy for a particle size of 40-nm GEM. For treatment of inhibitors in Fig. 3E–I, CHX, 2-DG, LatA, TBZ, CBZ, and Torin1 were added to cell suspensions at final concentrations of 100 μg/mL, 10 mg/mL, 10 μg/mL, 50 μg/mL, 60 μg/mL and 15 μg/mL, respectively.

For imaging of cells expressing μNS, the fluorescence images were recorded every 500 milliseconds for 2 min with the streaming mode. As in 40-nm GEM, the particles were tracked with the Mosaic suite using the following parameters: radius = 3; cutoff = 0; per/abs, variable; link range = 1; and maximum displacement = 5 pixels, assuming Brownian dynamics. Particles with tracks shorter than 10 frames were excluded from further analysis.

### Particle trajectory analysis

For individual trajectories of GEM and μNS particles, we computed the time-averaged mean-squared displacement (MSD) curves as in previous studies (7, 12). To compute the effective diffusion coefficient of individual particles, we truncated the MSD curves to the first 10 points, assumed that the linear relationship as

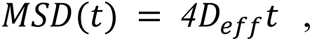

and used the least squares fitting. In order to estimate the power exponent of anomalous diffusion *α*, we averaged the individual MSDs for each condition. The ensemble-averaged MSDs were fitted to the following equation:

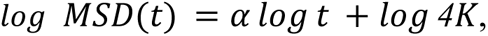

where *K* is the apparent diffusion coefficient in units of *μm^2^*/*S*^*α*^. In doing so, we used the ensemble-averaged MSDs from 100 ms to 1000 ms for the GEMs and from 1000 ms to 5500 ms for the μNS. In the figure legends, “n” indicates the number of trajectories, which was used to compute the effective diffusion coefficients.

### Quantification of trehalose amount

The amount of trehalose was measured by using a Trehalose Assay kit (K-TREH; Megazyme) with reference to the manufacturer’s instructions and the previously reported method (23). To compare the trehalose amount in vegetative cells and spores, cells grown exponentially in YEA medium were used as vegetative cells and spores were purified before use. Cell counts were measured by using Moxi GO II (ORFLO), and 1.0 × 10^8^ cells and 1.0 × 10^7^ cells were used as samples for the vegetative cells and spores, respectively. The cell volume was also measured using the same device (vegetative cells: about 30–40 fL; spores: about 10–12 fL). Cells were washed with 1 mL of YEA-glucose medium, and then supplemented with 100 μL of DDW heated at 95°C and vortexed vigorously. After incubation at 95°C for 15 min, the samples were centrifuged and about 100 μL of the supernatant was collected in new tubes. The pellets were treated with another 100 μL of DDW heated at 95°C and vortexed at room temperature for 5 min. After centrifugation, about 100 μL of the supernatant was collected in the tubes, and a total of about 200 μL of the cell extracts was obtained. Samples were measured on the day of extraction and stored at 4°C before use. 20 μL of each sample was measured in technical triplicate using a 96-well microplate (Thermo Fisher Scientific, #167008), and each measurement was performed in biological triplicate. Absorbance was measured at 340 nm using a Multiskan FC (Thermo Fisher Scientific). For absolute quantification of the trehalose amount, trehalose solutions (1.25, 2.5, 5.0, 10, 20 μg of trehalose) were also measured to create a calibration curve. To calculate the trehalose concentration, cell counts and volumes were corrected. To measure the intracellular trehalose amount during germination, 1.0 × 10^7^ cells were washed twice with DDW, suspended in YEA-glucose medium, and induced to germinate by adding glucose to a final concentration of 2%. At each time point during germination, cells were collected and the trehalose amount was quantified as described above.

### FCS measurement in fission yeast cells and analysis

FCS data were obtained using a Leica SP8 Falcon confocal microscope equipped with an objective lens, HC PL APO 63×/1.20 W motCORR CS2, and analyzed on Leica software essentially as described previously (76, 86, 87). Fission yeast cells expressing mNG were measured with 488 nm excitation and 500–600 nm emission. The structural parameter and effective confocal volume were calibrated using 500 nM Rhodamine 6G (TCI, R0039) in DDW based on the result that the diffusion constant of Rhodamine 6G in DDW is 414 µm^2^/s at room temperature (88). The Rhodamine 6G solution was measured with 488 nm excitation and 500–600 nm emission. The structural parameter and the effective confocal volume were estimated as 1.88 and 0.88 fL, respectively.

The time-series data of fluorescence fluctuations were obtained for 30 seconds, corrected by the photobleach correction algorithm of the Leica FCS analysis software, and subjected to the calculation of the auto-correlation function G(τ) of the Leica FCS analysis software. The calculated auto-correlation functions were fitted with the equation for the single-component normal diffusion and triplet model on Leica FCS analysis software, and effective diffusion coefficients of mNG were estimated.

### FRAP measurement in fission yeast cells and analysis

FRAP data were obtained using an FV3000 laser-scanning confocal microscope (Olympus) equipped with a UPLXAPO 60× objective lens, (NA = 1.42, WD = 0.15 mm; Olympus). Fission yeast cells expressing mNG were measured with 488 nm excitation and 500–540 nm emission. Both in vegetative cells and spores, about half of the cell area was photobleached and subsequent fluorescence recovery was observed. For photobleaching of cells, we used 100% laser intensity for 19 ms in vegetative cells and 40% laser intensity for 12 ms in spores. Fluorescence was recorded at an image acquisition rate of 50–100 ms, and a total of 150 images were acquired. Light stimulation was applied after 20 frames.

We estimated the effective diffusion coefficient by matching the observed FRAP data with the simulated data as explained below. To detect the cellular region of each cell, we averaged 20 pre-bleach frames and applied Otsu’s thresholding to binarize the time-averaged image. On the detected cellular regions with the no-flux boundary condition, we numerically simulated the spatio-temporal pattern of probe intensity, *c*(*x*, *y*, *t*), by using the diffusion equation

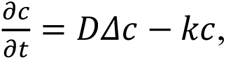

where *D* is the diffusion coefficient and *k* is the decay rate. We included the decay term to consider the observational photobleaching after light stimulation. Prior to the numerical simulation, the decay rate was estimated by linear regression on each time course of total fluorescent intensity within the cellular region. Then, we adopted the forward-time centered-space discretization (*Δt* = *0*.*005* [Frame], *Δx* = *1* [Pixel]) and numerically integrated the diffusion-decay model with different values of *D*. Using the same regions of interest (ROIs) as used for the experimental FRAP time courses *F*_*exp*_, we computed the time courses of mean intensity *I*(t) from the simulated images and rescaled them to obtain the simulated FRAP time course *F*_*Sim*_ as

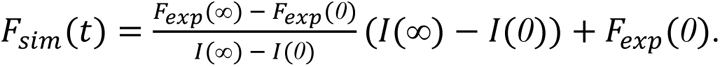

Finally, the optimal value of *D* for each cell was determined by grid search to minimize the mean-squared error between *F*_*Sim*_ and *F*_*exp*_, i.e.,

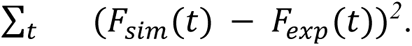

### Images and data analysis

All fluorescence imaging data were analyzed and quantified using Fiji/Image J (https://fiji.sc/). For all images, the background was subtracted using the rolling-ball method (rolling ball radius, 50.0 pixels). For the quantification of signal intensity in fission yeast cells, appropriate ROIs were manually selected, and mean intensities in ROIs were measured. To detect cells expressing mNG in Fig. 3C–D, mNG signals were directly segmented using Stardist (89, 90) and used as a proxy for cell area. Among the movie series (101 images) of 40-nm GEM particles, the first image was shown as a representative image. Data visualization and graph creation were performed using Python 3.10 with the Numpy 1.21.3, Pandas 1.3.4, Matplotlib 3.4.3, and Seaborn 0.11.2 modules.

## Acknowledgments

We thank all members of the Aoki Laboratory for their helpful discussions and assistance. We also thank members of the Tokai Tor Conference (ToToCo) for the general discussion. Some fission yeast strains were provided by the National Bio-Resource Project (NBRP), Japan, and others were a kind gift from Dr Jun-ichi Nakayama. K.A. was supported by JSPS KAKENHI grants (nos. 18H02444, 19H05798, and 22H02625). Y.G. was supported by a JST, ACT-X grant (no. JPMJAX22B8), by JSPS KAKENHI grants (nos.19K16050 and 22K15110), by a Jigami Yoshifumi Memorial Research grant, and by a Sumitomo Research grant. K.S. was supported by a JSPS KAKENHI grant (no. 22J10844) and The Graduate University for Advanced Studies, SOKENDAI (SOKENDAI Student Dispatch Program).

## Competing Interests

No competing interests declared.

## Author contributions

K.S., Y.G. and K.A. designed the research; K.S. and Y.G. performed the experiments; K.S., Y.G. and Y.K. analyzed the data; K.S., Y.G., Y.K. and K.A. wrote the manuscript; all authors approved the final manuscript.

## Notes

### Competing Interest Statement

The authors have declared no competing interest.

