## Supplementary Information for "Cytoplasmic fluidization triggers breaking spore dormancy in fission yeast"

fission yeast, germination, cytoplasmic fluidity, cAMP-PKA pathway, trehalose

### Supplementary Figures

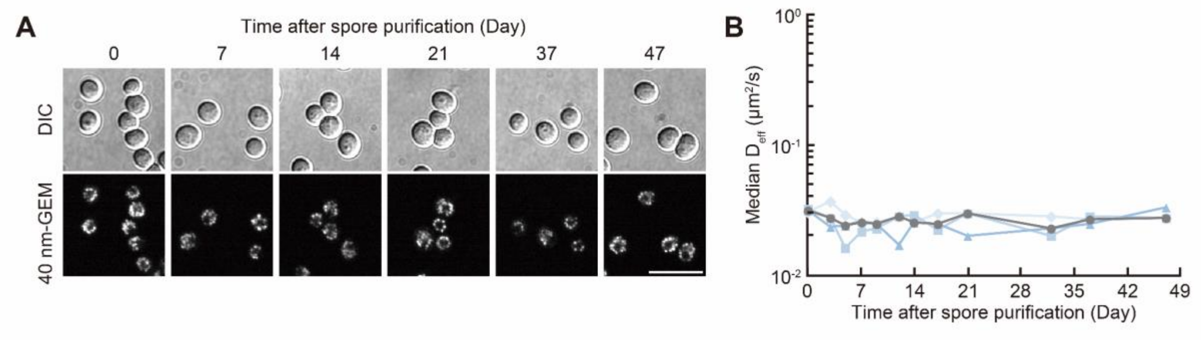


**Fig. S1. The mobility of 40 nm particles does not change during the storage of spores at 4°C.**

(A) Representative DIC (upper) and confocal fluorescence (lower) images of fission yeast spores expressing 40-nm GEMs after purification. Purified spores were stored at 4°C for the indicated days and imaged. Scale bar, 10 µm.

(B) Median $D_{\text{eff}}$ of 40-nm GEMs in spores stored at 4°C for the indicated periods (0, 3, 5, 7, 9, 12, 14, 17, 21, 32, 37, 47 days). Blue lines represent the median $D_{\text{eff}}$ for each of the three independent experiments, and a gray line indicates their mean (n > 850 for each condition).

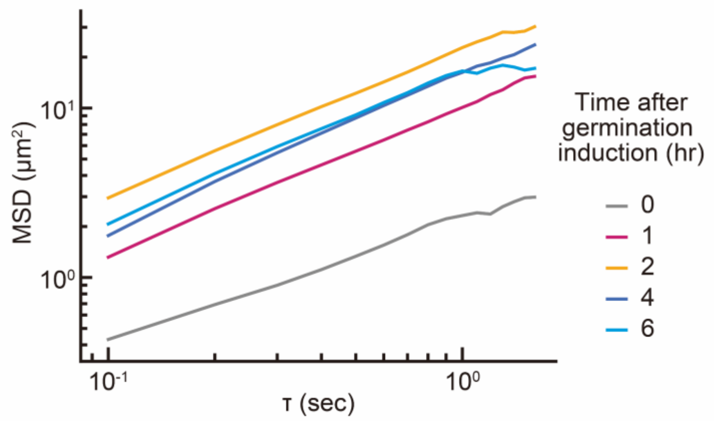


**Fig. S2. The mobility of 40 nm particles is increased during the initial stage of spore germination.**

Ensemble-averaged MSD curves of 40-nm GEMs during germination (0 min, n = 801; 60 min, n = 925; 120 min, n = 1023; 240 min, n = 533; 360 min, n = 416).

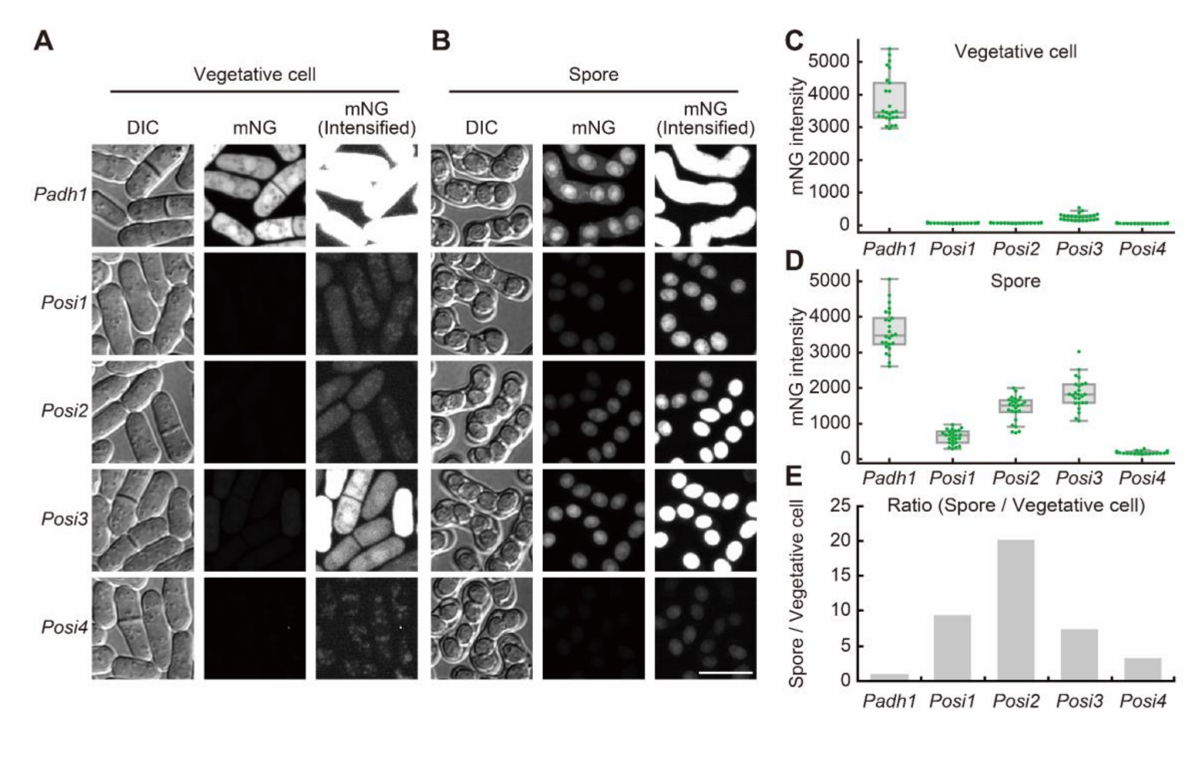


**Fig. S3. Development of a spore-specific gene expression system in fission yeast.**

(A and B) Representative DIC (left) and confocal fluorescence images (middle and right) of vegetative cells (A) and spores (B) expressing mNeonGreen (mNG) under the constitutive promoter *Padh1*, or the spore-inducible promoters *Posi1-4*. Because the expression levels of mNG under *Posi1-4* were much weaker than those of *Padh1*, the intensified fluorescence images are shown in the right column. Scale bar, 10 μm.

(C and D) Quantification of the mNG fluorescence intensities in vegetative cells (C) and spores (D). Each dot represents mNG fluorescence from a single cell with a boxplot, in which the box shows the quartiles of data with the whiskers denoting the minimum and maximum except for the outliers detected by 1.5 times the interquartile range (n = 25 cells).

(E) Ratio of mean mNG fluorescence intensity of spores to that of vegetative cells.

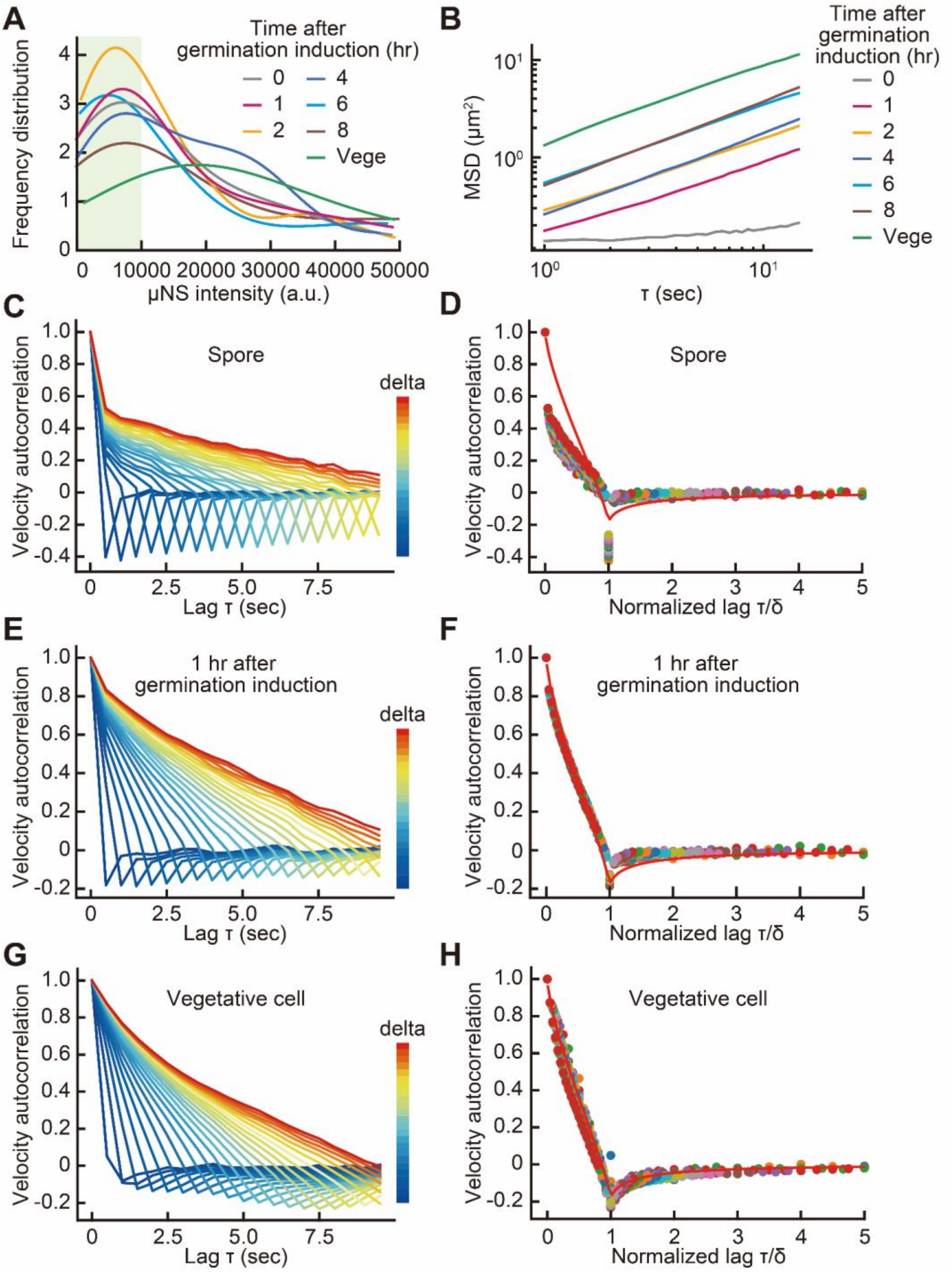


**Fig. S4. Analysis of μNS trajectories.**

(A) Distribution of fluorescence intensity of μNS particles in spores at the indicated time after germination induction and vegetative cells (Vege) (n > 90 trajectories for all conditions).

(B) Ensemble-averaged MSD curves.

(C-H) Normalized velocity autocorrelation function (VAF) $c^{\delta}(\tau)/c^{\delta}(0)$, where $c^{\delta}(\tau) = \langle v(t+\tau)\cdot v(t)\rangle$, $v(t) = (r(t+\delta) - r(t))/\delta$, and $r(t)$ is a coordinate vector at Time $= t$ in a trajectory. For comparison, we chose the trajectories in the spore cytoplasm (C-D), the cytoplasm 1 hr after germination induction (E-F), and the log-phase cytoplasm (G-H). Ensemble-averaged VAFs are plotted for lag time $\tau$ (C, E, G) and normalized lag time $\tau/\delta$ (D, F, H). Red solid lines in (D, F, H) represent the theoretical VAF for fractional Langevin motion (anomalous exponent = 0.7), which was computed from Equation (1) in (1).

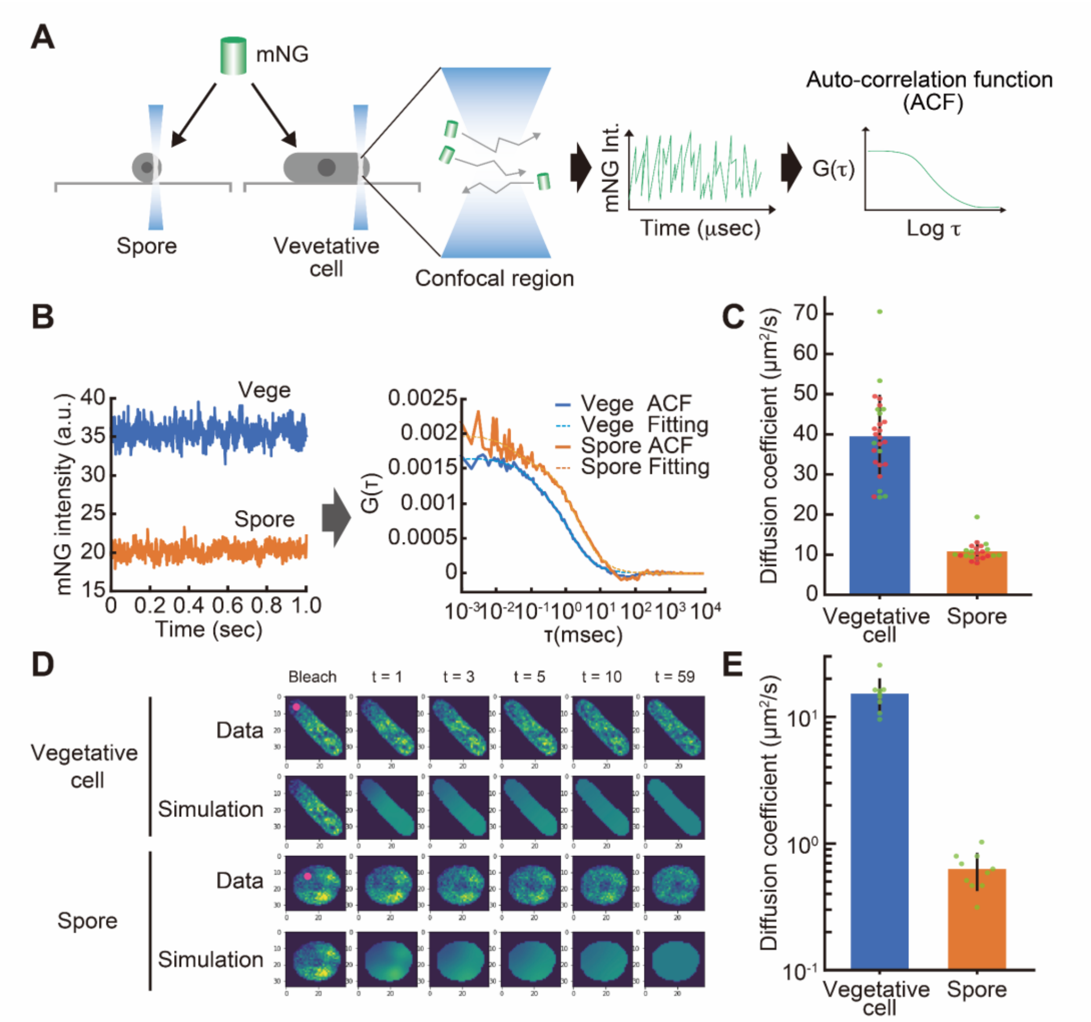


**Fig. S5. Quantification of the mNG diffusion coefficient in the spore cytoplasm by FCS and FRAP.**

(A) A schematic illustration of fluorescence correlation spectroscopy (FCS) analysis in fission yeast cells. mNG fluorescence fluctuations in a confocal volume were measured in vegetative cells and spores, and then auto-correlation functions G(τ) were calculated from the time-series data. Diffusion coefficients of mNG were computed according to the procedures described in Materials and Methods.

(B) Representative FCS data obtained from cells expressing mNG. The left graph represents the raw data of mNG fluorescence fluctuation in vegetative cells and spores. The right graph shows the calculated autocorrelation functions (ACF, lines) and the fitted curves (Fitting, dashed lines) (for details see Materials and Methods).

(C) Quantification of diffusion coefficients for mNG in vegetative cells and spores by FCS. Diffusion coefficients for more than 10 cells were calculated from two independent experiments and plotted with different colored dots (green or red). The bar plot indicates the mean values with error bars (SD).

(D) Representative images of fluorescence recovery after photobleaching (FRAP) analysis in fission yeast cells (for details see Materials and Methods).

(E) Quantification of diffusion coefficients for mNG in vegetative cells (n = 9 cells) and spores (n = 10 cells) by FRAP analysis.

**Movie S1. The mobility of 40 nm particles in vegetative cells and spores of fission yeast.** Representative differential interference contrast (DIC) (upper) and spinning disk confocal fluorescence images (lower) of vegetative cells (left) and spores (right) of fission yeast expressing 40-nm GEMs. The fluorescence images were taken every 100 milliseconds, and the movie playback speed is set to 20 frames per second. The timestamp shows time in seconds. Scale bar, 10 μm.

**Movie S2. The mobility of 40 nm particles during germination in fission yeast.** Representative differential interference contrast (DIC) (upper) and spinning disk confocal fluorescence images (lower) of fission yeast expressing 40-nm GEMs at the indicated time after germination initiation. The fluorescence images were taken every 100 milliseconds, and the movie playback speed is set to 20 frames per second. The timestamp shows time in seconds. Scale bar, 10 μm.

**Table S1. Plasmid list**

| **Plasmid name** | **Description** | **Source** | **Benchling Link** |
|---|---|---|---|
| pMNAT2LA21-40nm-GEM_sp_opt | 40-nm GEMs optimized for *S. pombe* codon usage | this study | https://benchling.com/s/seq-O4wGxsTRYMRIaGhT08sC?m=slm-DksK0Fql7vV3NaAZrC47 |
| pMNATZA1-spmNeonGreen | | K. Sakai et al., 2021 | https://benchling.com/s/seq-8QqFWbwYIEF8y0PeV022?m=slm-AqJJ200oBOTLichEvFb7 |
| pHBCO1-HA-spmNeonGreen | | this study | https://benchling.com/s/seq-PmMNBPWi2rhdjj6aDS2X?m=slm-fgSfoDqEYQpszprFlisB |
| pHBCO2-HA-spmNeonGreen | | this study | https://benchling.com/s/seq-K9M0fzPKUL8o5lJmNNVW?m=slm-pnnt1t5deDZTQ4J43O3I |
| pHBCO3-HA-spmNeonGreen | | this study | https://benchling.com/s/seq-DdD2RR1IAVNSpp139v07?m=slm-qT3NV619uAU80C7RVx00 |
| pHBCO4-HA-spmNeonGreen | | this study | https://benchling.com/s/seq-9zNLv2iwVBmo4DgerOww?m=slm-dpjV6HCdFGBkJdEohC9K |
| pHBCO2-ntp1 | | this study | https://benchling.com/s/seq-jvtAd53Mw9VgFl4YFw1X?m=slm-oXvW89tp4XNoEnRFF3wo |
| pHBCA1-HA-spmNeonGreen-new | | this study | https://benchling.com/s/seq-Hr3wZ7ZFqDyLcwXsUTRo |
| pMNATZA1-GFP-µNS | | this study | https://benchling.com/s/seq-B1IjgWLExt3H0W0Y9PlS |

**Table S2.** ***Schizosaccharomyces pombe*** **strain list**

| Strain name | Genotype | Fig. | Source |
|---|---|---|---|
| L972 | h- | | NBRP |
| L975 | h+ | | NBRP |
| L968 | h90 | Fig. 2B, 2C, Fig. 4A, 4B, 4C, 4G | NBRP |
| SK445 | h90 2L::Padh21-40nm-GEM<<nat | Fig. 1B, 1C, 1D, 1E, 1F, 1G, Fig. 2D, Fig. 3A, 3B, 3E, 3F, 3G, 3H, 3I, Fig. 4B, 4D, 4E, 4F, 4I, 4J, 4K, Fig. 5D, Fig. S1, Fig. S2A, S2B | this study, L968 |
| TN366 | h+ ade6-M216 leu1-32 | | J. Nakayama Lab |
| SP976 | h90 ade6-M210 leu1-32 ura4-D18 | | J. Nakayama Lab |
| YG008 | h90 ade6-M210 leu1-32 | | this study, TN366 × SP976 |
| SK009 | h90 ade6-M210 leu1-32 pka1::hyg | Fig. 2B, 2C | this study, YG008 |
| SK012 | h90 ade6-M210 leu1-32 cyr1::nat | Fig. 2B, 2C | this study, YG008 |
| SK028 | h90 cyr1::kan | | this study, L968 |
| SK520 | h90 git3::kan | | this study, L968 |
| SK533 | h90 pka1::kan | Fig. 4C, 4G | this study, L968 |
| SK570 | h90 cyr1::kan 2L::Padh21-40nm-GEM<<nat | Fig. 2D | this study, SK028 |
| SK571 | h90 git3::kan 2L::Padh21-40nm-GEM<<nat | Fig. 2D | this study, SK520 |
| SK577 | h90 pka1::kan 2L::Padh21-40nm-GEM<<nat | Fig. 2D | this study, SK533 |
| SK037 | h90 z::Padh1-spmNeonGreen<<nat | Fig. 3C, 3D, Fig. 5D, Fig. S3A, S3B, S3C, S3D, S3E, Fig. S5B, S5C, S5D, S5E | this study, L968 |
| SK528 | h90 ntp1::kan | Fig. 4C, 4G | this study, L968 |
| SK575 | h90 ntp1::kan 2L::Padh21-40nm-GEM<<nat | Fig. 4D, 4E, 4F | this study, SK528 |
| SK590 | h90 gpa2::kan | | this study, L968 |
| SK612 | h90 gpa2::kan 2L::Padh21-40nm-GEM<<nat | Fig. 4I, 4J, 4K | this study, SK590 |
| SK668 | h90 gpa2::kan 2L::Padh21-40nm-GEM<<nat c::Posi2-ntp1<<hyg | Fig. 4I, 4J, 4K | this study, SK612 |
| YG430 | h90 c::Posi1-HA-spmNeonGreen<<hyg | Fig. S3A, S3B, S3C, S3D, S3E | this study, L968 |
| YG431 | h90 c::Posi2-HA-spmNeonGreen<<hyg | Fig. S3A, S3B, S3C, | this study, L968 |

| | | | |
|---|---|---|---|
| | | S3D, S3E | |
| YG432 | h90 c::Posi3-HA-spmNeonGreen<<hyg | Fig. S3A, S3B, S3C, S3D, S3E | this study, L968 |
| YG433 | h90 c::Posi4-HA-spmNeonGreen<<hyg | Fig. S3A, S3B, S3C, S3D, S3E | this study, L968 |
| SK425 | h90 z::Padh1-GFP-uNS<<nat | Fig. 5A, 5B, 5C, 5D, Fig. S4A, S4B, S4C, S4D, S4E, S4F, S4G, S4H | this study, L968 |
